# Visual detection of cryptic displays in jumping spiders

**DOI:** 10.64898/2026.04.03.716102

**Authors:** Massimo De Agrò, Francesca Lo Bello, Peter Neri, Giorgio Vallortigara

## Abstract

Most animals can segment camouflaged targets via static cues, such as luminance or contrast differences against a background. However, targets can be cryptic and merge with background clutter, making static cues unavailable until the camouflaged object starts moving: the target can then be segmented by identifying spatially coherent luminance changes over time. Jumping spiders represent a unique model for the study of spatiotemporal segmentation, as they process moving signals and static figures separately through distinct pathways associated with different pairs of eyes: the secondary eyes detect moving objects and trigger full body pivot responses, bringing the target within focused view of the principal eyes for subsequent figure processing. While secondary eyes can be selective for specific stimulus characteristics that cause the animal to pivot, it remains unclear whether their selectivity requires full motion segmentation or relies on simpler stimulus characteristics. Here, we tested whether the pivoting response triggered by secondary eyes is specifically linked to motion segmentation, or whether it is also elicited by less specific stimulus characteristics. We presented jumping spiders with several moving and non-moving stimuli, including clearly delineated objects, cryptic objects visible only through motion, and non-segmentable spatio-temporal luminance patterns. Based on the measured responses, we developed a computational model that captured all salient features of the pivoting behavior produced by the animal. We found that jumping spiders do not require a discrete, segmented object to trigger a pivoting response. Instead, they attend to luminance patterns that change across both time and space, while disregarding changes that repeat over the same spatial location. This mechanism constitutes an effective solution in terms of neural simplification: it offloads the task of complex motion segmentation from the secondary eyes, thus maximizing the efficiency of visual modularity—a unique evolutionary solution to the problem of brain miniaturization.

## 1 Introduction

Animals are under constant pressure to identify relevant objects against cluttered backgrounds, a task that requires figure-ground segmentation. This ability has been of interest to scientists at least since the time of Gestalt psychology [1]. The rules of figure-ground segmentation have been mostly discussed in relation to static images, in which target objects can be isolated from their background via overall differences in luminance or the presence of visible edges. Yet, sometimes the luminance difference between an object and its background is blurred (often intentionally, as is the case for cryptic organisms [2]), making the segmentation process unreliable. However, as soon as an object starts moving, its edges become immediately evident to us, popping out of the background as if they were the only salient elements against a black canvas. This phenomenon is driven by perceptual grouping of luminance variations, whereby the percept of a whole object is supported by spatiotemporal coherence [3, 4, 5].

Motion per se is, however, insufficient for reliably detecting objects: not every luminance change over time is caused by a segmentable agent—on the contrary, it can be generated by self-motion, global light-source occlusions, or moving shadows. In order to effectively detect objects in the environment, visual systems must be able to separate the different potential sources of motion signals and further elaborate motion information using the rules of static perception [6].

Jumping spiders have lately gained attention in the area of motion-based and figure-based segmentation, due to the unique manner in which their visual system processes spatiotemporal information [7, 8]. Spiders of the jumping family possess four different pairs of eyes, each specialized for a different visual task. The biggest, front-facing pair, called principal or anterior medial eyes (AME), carry the highest visual acuity at the cost of a narrow field of view [9, 10], and are believed to be specialized for figure recognition [10]. The other three pairs, collectively named secondary eyes, encompass anterior lateral (ALE), posterior medial (PME), and posterior lateral (PLE) eyes. They carry lower visual acuity compared with the AME, but collectively equip the spider with nearly 360° of field of view [11, 12, 13], and are responsible for motion detection [12]. Regardless of which target they detect, the secondary and primary eyes of jumping spiders work synergistically, as evidenced by the behavior produced by the animal: as soon as an object moves across the visual field of the secondary eyes, the spider performs a sudden whole-body pivoting movement towards it [12, 14, 15], in order to bring the object within direct view of the AME. Thereafter, the retinas of the AME engage in a semi-stereotyped “scanning” movement across the target [10], until the figure is recognized [16, 17, 18].

Crucially, jumping spiders do not produce pivots for any perceived change in luminance. On the contrary, they can be quite selective with respect to which stimuli may trigger a pivoting response [19, 20], including the ability to select one target among many [21, 13, 22]. Furthermore, secondary eyes are not all equally selective, but seem instead to present various levels of specialization. For example, ALE can finely discriminate motion patterns and infer target structure [13, 22, 23], and are also responsible for directing the movements of AME [14, 24, 25]. All evidence regarding the pivoting behavior of jumping spiders points towards complex processing, rather than simple luminance-change detectors. To summarize (and inevitably simplify) current consensus in the literature, it seems that pivoting responses are only produced when an actual object is detected, suggesting that secondary eyes perform the task of figure-ground segmentation directly from motion, and then pass this information onto the AME for further inspection of already pre-isolated objects for recognition [19, 13, 22, 24]. Yet, no direct test of this hypothesis has ever been performed, leaving many open questions regarding the process governing pivot production and motion detection in this model species.

In this study, we tested the rules governing motion-based object segmentation and subsequent body re-orientation in jumping spiders. In order to do so, we placed individual animals inside a virtual reality setup and presented them with different stimuli displayed on computer monitors covering the visual field of secondary eyes. The stimuli could present either a segmentable object or no object at all but still be characterized by luminance changes. Moreover, some of the presented objects could be segmented via luminance cues, while others required specific reliance on motion cues. This set of stimuli was designed with the goal of disentangling different cues and identifying what stimulus characteristics may be regarded as constitutive of an “object” by the visual system of jumping spiders, and hence capable of triggering a pivot response. Subsequently, we repeated our measurements by presenting stimuli only in subportrions of the spider’s visual field, in order to test whether the same perceptual rules triggering pivots are shared across eyes. Finally, we developed a computational model capable of comprehensively capturing the observed behavior.

## 2 Materials and Methods

### 2.1 Subjects

A total of 100 individuals of *Menemerus semilimbatus* were used across the three experiments (31 for the first, 36 for the second, and 33 for the third). Specimens were collected in the wild between March and May 2024. Collection and housing procedures followed established protocols [22]. After capture, each spider was transported to the laboratory and subsequently transferred to a ventilated plastic container (16 × 8 × 5.5 cm) equipped with a water source and a cardboard shelter. Spiders were maintained under controlled lighting conditions with a 12h light/dark cycle. On the day following capture, each individual underwent a magnet attachment procedure following the methodology described in [22]. The spider was gently immobilized between a sponge and a latex membrane, which had an opening aligned with the cephalothorax. A 1 × 1 mm cylindrical magnet was then affixed to the cephalothorax using a resin, which was rapidly polymerized under a UV lamp. After magnet attachment, each spider remained in its container for 24 hours before the experiment began, allowing for habituation to the presence of the magnet. At the end of each experiment, the magnet was mechanically removed and the spider released back in nature.

### 2.2 Experimental Procedure

For a full description of the experimental apparatus, please see [22]. Briefly here, the spider was attached to the magnetic end of a 6-axis manipulator and positioned such that its legs came into contact with a 38-mm polystyrene sphere, which was suspended by a stream of compressed air to make it frictionless. Under this setup, the spider could not alter its position/orientation, but was free to move its legs and to impress its desired direction to the sphere. The latter was recorded at 320 × 240 pix, 120 fps, and its movement reconstructed through the software FicTrac [26]. We placed a computer monitor in front of the spider at a viewing distance of 15.7 cm, where stimuli were presented at 1080p (1920 × 1080 pix) and 30 fps. For experiment 3 we used two monitors, one to the left and one to the right of the spider, forming a 40° angle between them. This arrangement was chosen to permit selective independent stimulation of ALE and PLE. The distance between the monitor centers and the spider was kept at 15.7 cm, so as to retain the same angular size of displayed stimuli. At this distance, 1° of visual angle approximately equated 10 pixels. At the start of each trial, monitor(s) displayed a random checker pattern of white and black squares (in equal number), each measuring 0.4°. At the beginning of each trial (consisting of multiple stimulus presentations), a 210-seconds habituation period was provided, during which only the background (checker pattern) was displayed and no stimulus was shown. After this period, a total of 50 stimuli were presented, each stimulus lasting 1.33 seconds with an inter-stimulus interval of 15 seconds.

#### Stimuli

We used for the experiment a total of 8 different stimuli, with 5 of these used for experiment 1 and 6 for experiments 2 and 3. See Figure 1A–G for a spatio-temporal plot of the stimuli and Video 1 for a full depitction.

**Figure 1:**
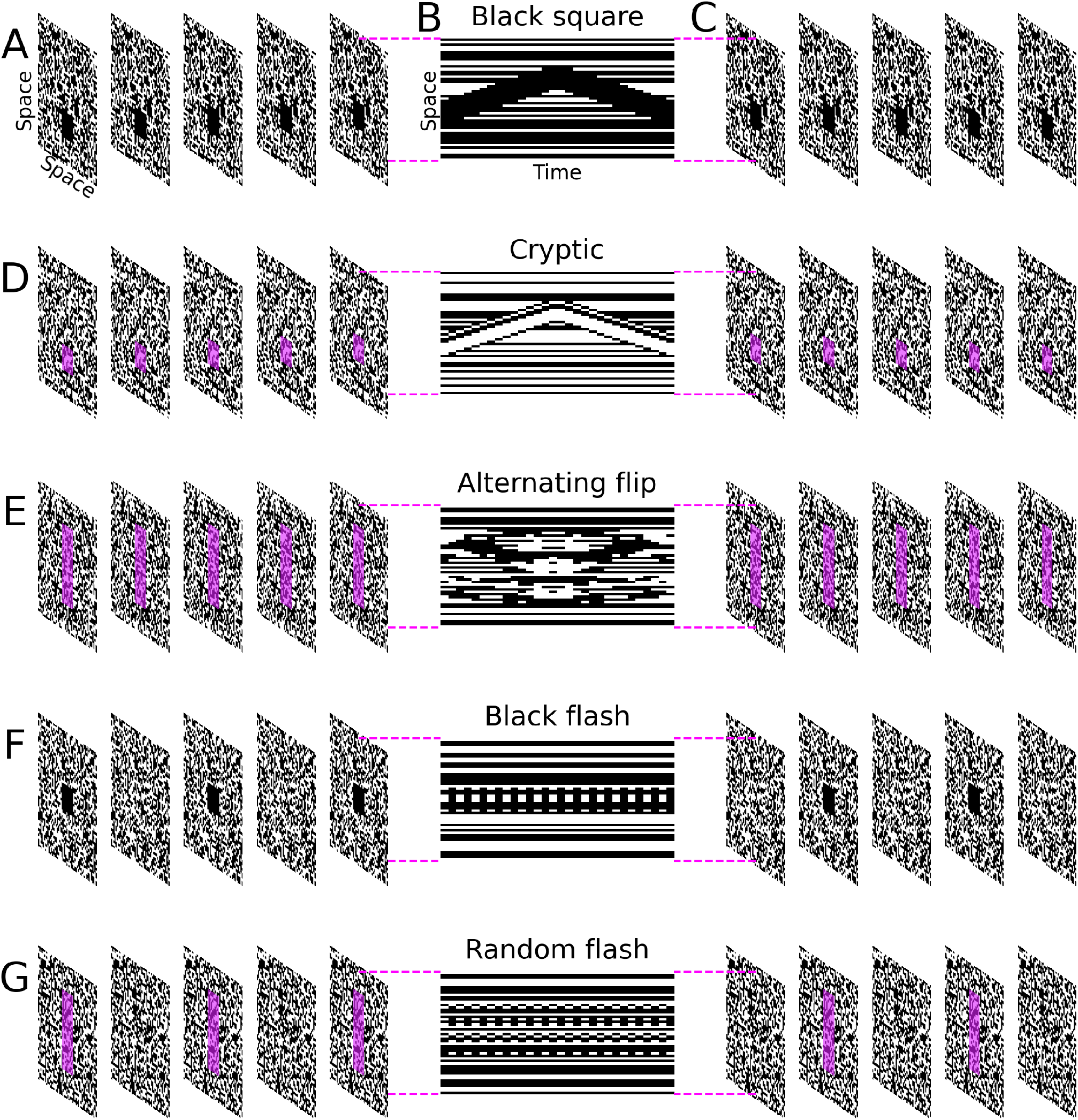
Stimulus specification. **A** shows first five frames of the “Black square” stimulus in perspective view. **B** shows subsequent frames as space-time diagram in which the intensity along the space dimension reflects the intensity of pixels traversing the vertical meridian of each frame. **C** shows final five frames in perspective view. The “Gray square” and “White square” stimulus are identical, except for the intensity of the moving square region. **D** shows the “Cryptic” stimulus using the same plotting conventions. For frames in perspective view, the moving region is indicated by magenta shading. **E** shows the “Alternating flip” stimulus, with the manipulated region highlighted by magenta shading. **F** shows the “Black flash” stimulus. The “White flash” stimulus was identical except for the intensity of the flashed square region. **G** shows the “Random flash stimulus, with the manipulated region (replaced by a pixel-noise sample that differed from the background) highlighted by magenta shading. See also Video 1 for the stimuli in movie form.

The “Black square” stimulus (Figure 1A–C) consisted of a 4 × 4° black square presented at level height with the spider’s head. After appearance, the square moved upwards for a distance of 10° in 0.66 seconds. It then reversed direction and returned to its original location, after which it disappeared.

The “Grey square” and “White square” (Figure 1A–C, but with different luminance) stimuli were identical except for the square taking on a different luminance level, as indicated by their denominations. Because of its homogeneous pattern, the square element in all these stimuli could be easily segmented from the background on individual frames (Figure 1A), without necessitating movement over multiple frames: it can be segmented using either shape or motion.

In the “Cryptic” stimulus (Figure 1D), the pattern within the square element consisted of white and black pixels that were statistically indistinguishable from the background, so that the square element could not be segmented on individual frames: the square element could be detected only while in motion, perfectly blending with the background if seen as a static snapshot.

In the “Alternating Flip” (Figure 1E) stimulus, the active area measured 4° × 10° (width × height), so as to cover the entire region spanned by the square element in the other stimuli as it traveled vertically during stimulus presentation. Within this area, we randomly selected a subset of squares equal in number to half the number of squares composing the Cryptic stimulus. On each frame, we flipped the polarity of the selected squares between black and white, so that the overall amount of luminance change per unit time remained identical to that associated with the Cryptic stimulus, while no translation was effected: in the Alternating Flip stimulus, there is no clearly segmentable object, whether within frames or across frames, except for an overall region of interest within which some elements vary their contrast. If jumping spiders selectively produce pivoting movements following detection of whole objects, they should not respond to the Alternating Flip stimulus.

In the “Black Flash” (Figure 1F), a 4° wide black square appeared on screen, to then disappear again in the next frame. The stimulus continued appearing and disappearing in the same location for a total of 1.33 seconds. From a previous experiment, we already know that spiders should not respond by pivoting when presented with this stimulus [19]. Note, however, that they do perceive it, as its presentation changes the spiders’ subsequent reaction to other experimental stimuli [19]. Crucially, this flashing stimulus does present a clear segmentable object, but it’s characterized by no spatial changes in luminance, only temporal. This is in contrast with the Alternating Flip stimulus, where there is no segmentable object, but the luminance changes in space and in time.

The “White Flash” (Figure 1F, but with flipped luminance) stimulus was identical to the previous one, but with a white square rather than black.

The “Noise Flash” (Figure 1G) stimulus was constructed starting from the same logic of the Alternating Flip one. In a 4 by 10 degrees area, we selected a sub-portion of the white and black squares, equal to the number of squares flipping each frame for the black or white translating stimuli. As per the Alternating Flip stimulus, these squares flipped from black to white or from white to black. In the next frame, the same exact squares flipped again, continuing this alternation for a total of 1.33 seconds. As such, differently from the Alternating Flip stimulus, in the Noise Flash the luminance change did not spatially vary. Moreover, differently from the Black and White flash, no segmentable object was visible.

**Experiment 1** In experiment 1, we presented five different stimuli, specifically the “Black square”, “Grey square”, “White square”, “Cryptic” and “Alternating flip”. Each stimulus was presented 10 times, randomly shuffled across the full trial. Stimuli could appear on either side of the monitor (left *vs*. right) at *±* 40° with respect to the vertical meridian of the spider’s line of sight. This mode of presentation placed stimuli well outside the visual field of the AME [11] but well inside that of the ALE [13].

**Experiment 2** In experiment 2, the monitor positioning and stimuli locations remained the same. Moreover, we presented again the Black, the Cryptic and the Alternating Flip stimuli, but we added three new stimuli, specifically the Black flash, White flash and Noise flash (Figure 1F–G, Video 1). This was prompted by the unexpected responses that the spiders produced to the Alternating Flip stimulus (see results), where we expected no reaction due to the absence of a segmentable object, but only unstructured luminance change.

**Experiment 3** In experiment 3, we used again the same exact stimuli of experiment 2. However, we added a computer monitor to the setup, and moved the two screens to be at each side of the spider, rather than in front. The two monitors formed an ideal 40° angle between them (without actually touching), with its centers aligned with the spider position and 157 mm afar. This was done to maintain the angular sizes of experiments 1 and 2. This monitor positioning allowed for the presentation of stimuli both at *±* 40° (and so visible to ALEs) and at *±* 120° (and so visible to PLEs) [13]. To avoid unwanted influences, each subject was assigned to either the ALEs condition or the PLEs condition, and as such only saw stimuli in the ALEs fields or the PLEs fields.

### 2.3 Scoring

The scoring procedure was the same as in [22]. In brief, the X, Y and Z rotations of the spherical treadmill as extracted by fictrac [26] were temporally aligned with the stimuli presentation. Thereafter, we selected 2 seconds long of the data starting from stimuli appearance. Here, we checked for the presence of rotations of the ball around its Z axis without synchronous movement in the X or Y (i.e., pure rotations around the vertical axis) in the direction of the stimulus position. In the presence of such rotation, we considered the spider to have reacted to the stimulus, ending up with a binomial variable. To control for false positives, we also extracted from the data 50 other 2-seconds long sections from the habituation phase, and as such were not aligned with any stimulus appearance. Here, we tested for the same presence of Z ball rotations, as to find a baseline probability of responses (i.e., no-stimulus control). During data analysis, we included this sections in the model, and tested the response rate of the spiders *vs*. this baseline. Absence of a statistical difference from this control was interpreted as the spiders not responding to the stimulus. The scoring was fully performed in Python 3.12 [27], using the packages Pandas [28], Numpy [29, 30] and Scipy [31, 32].

### 2.4 Statistical Analysis

All analyses were performed in R 4.4.2 [33]. After loading the data using the package data.table [34], for all three experiments we fitted a generalized linear mixed effect model based on a binomial error structure using the package glmmTMB [35]. We set as our dependent variable the spiders’ response (either pivoting towards the stimulus or not). As independent variables, for experiment 1 and 2 we only used the stimulus type (including the no-stimulus control). For experiment 3 instead we used the stimulus type and its location (either in the ALEs or the PLEs visual field). We used the subject identity as random intercept. After checking the goodness of fit with the package DHARMa [36], we analyzed the predictors effect with an analysis of deviance through the package Car [37]. Then, we performed post-hoc analysis on significant predictors using the package Emmeans [38], applying a Bonferroni correction. Lastly, we produced plots using both the package Ggplot2 [39] and the Python[27] library Matplotlib [40].

### 2.5 Computational model

We implemented a variant of the NLN cascade model classically used to capture visual processes in insect vision [41, 42].

#### Input specification

Each input stimulus can be represented in the form of a 3D array *I*(*x, y, t*): two dimensions of space *x* and *y* spanning each 2D frame, and the remaining dimension of time *t* spanning different frames (**Figure 1**). In our simulations, each element of *I* could take one of three values: 0 for dark, 0.5 for gray, and 1 for bright.

#### Early nonlinearity

We first subjected the input array to squaring, effectively implementing a threshold-quadratic nonlinearity (**Figure 3C**). This type of static expansive transduction is meant to emulate the known gamma-expansion nonlinearity of most computer display systems, which we confirmed for our own display (photometric measurements of black/gray/white on our monitor returned lux values of 0/36/164). However, it may also capture some aspects of biological photo-transduction [43].

#### Front-end filtering

The squared input *I*(*x, y, t*)^2^ was fully convolved (**Figure 3D**) with a 3D filter specified by seven frames (equivalent to an overall duration of 230 ms). The filter was constructed by multiplying a 2D difference-of-Gaussians (DOG) function with standard deviations (SDs) of 0.2/0.3 degrees (for positive/negative Gaussian blobs, the peak of the latter being set to twice the peak of the former) with a sinusoidal temporal impulse response spanning one peak-to-trough half-cycle over the 7 frames (**Figure 3E**). We normalized this filter to unit energy so that numerical values for model parameterization (see below) can be expressed in meaningful/reproducible units.

#### Late nonlinearity

The 3D array returned by front-end convolution was subjected to full-wave rectification via squaring (**Figure 3F**), effectively implementing energy extraction [44].

#### Addition of internal noise

Following the above pipeline, the resulting output still consisted of a 3D array (**Figure 3G**). Based on extensive experimental observation of spider behavior, we determined that spider typically initiated their turning behavior well before completion of stimulus presentation. We therefore focused on the first 500 ms of the output 3D array and summed energy within this time window (**Figure 3H**), resulting in scalar output *y*. This output was further perturbed by a noise source (**Figure 3I**) that simulates intrinsic behavioral variability via the addition (**Figure 3J**) of a scalar value from a Gaussian distribution with SD equal to *y*^2^ + *k*, where *k* = 1.2. In this specification, internal noise scales supralinearly with *y* [45] and incorporates a “dark noise” component *k* [46].

#### Conversion to binary turn/stay decision

When the noisy output exceeded a threshold of 1.8 (**Figure 3K**), the model produced a “turn” response (**Figure 3L**); when threshold was not reached, it produced a “stay” response (**Figure 3M**).

## 3 Results

Only the main results are reported here. For the complete analysis, raw data and python scripts, see [47]. For all three experiments, as a first step we selected subjects to be included in subsequent analysis. We already know from previous literature that the animals are capable of detecting the up-down moving black and white squares. As such, we decided to drop all subjects for which the response rate to this control stimulus was lower than their baseline activity (i.e., response rate to sections where no stimulus was visible on screen). We provided in SI2 also the analysis on the full set of spiders. For the most part the observed effects are identical, with the only difference being the absolute probability values being much lower due to the presence of non-responders.

### 3.1 Experiment 1

For experiment 1, 12 out of the 31 subjects (38.7%) responded to less than 10% of the control stimuli, and as such were dropped from the subsequent analysis. We did observe a general effect of stimulus type (GLMM ANODA, *χ*^2^=191.7, DF=5, p<0.0001). Specifically, the spiders showed a response level significantly different from their baseline activity to all of the stimuli (Figure 2. GLMM post-hoc, Bonferroni corrected. Black: odds ratio=5.38, SE=0.951, z=9.517, p<0.0001. Grey: odds ratio=4.492, SE=0.798, z=8.455, p<0.0001. White: odds ratio=0.469, SE=0.0898, z=3.957, p=0.0011. Alternating flip: odds ratio=0.133, SE=0.0236, z=11.364, p<0.0001. Cryptic: odds ratio=0.325, SE=0.0593, z=6.162, p<0.0001.). Moreover, we observed no significant difference in the response rate between the black, gray and alternating flip. Moreover, we observed no difference between the cryptic and the white stimulus. The response rate to the white stimulus resulted significantly lower than all the three former stimuli (Figure 2. Black *vs*. white: odds ratio=2.522, SE=0.567, z=4.11, p=0.0006. Grey *vs*. white: odds ratio=2.105, SE=0.475, z=3.297, p=0.0147. Alternating flip *vs*. white: odds ratio=3.526, SE=0.795, z=5.588, p<0.0001.), while the cryptic stimulus resulted significantly lower only from the alternating flip stimulus (Figure 2. Black *vs*. cryptic: odds ratio=1.747, SE=0.379, z=2.57, p=0.1523. Grey *vs*. cryptic: odds ratio=1.459, SE=0.318, z=1.732, p=1. Alternating flip *vs*. cryptic: odds ratio=2.443, SE=0.531, z=4.107, p=0.0006.). It is notable that the response rate to the alternating flip is significantly higher than to cryptic, as we would have expected no response. The alternating flip should present in fact no segmentable object, differently from all other stimuli.

**Figure 2:**
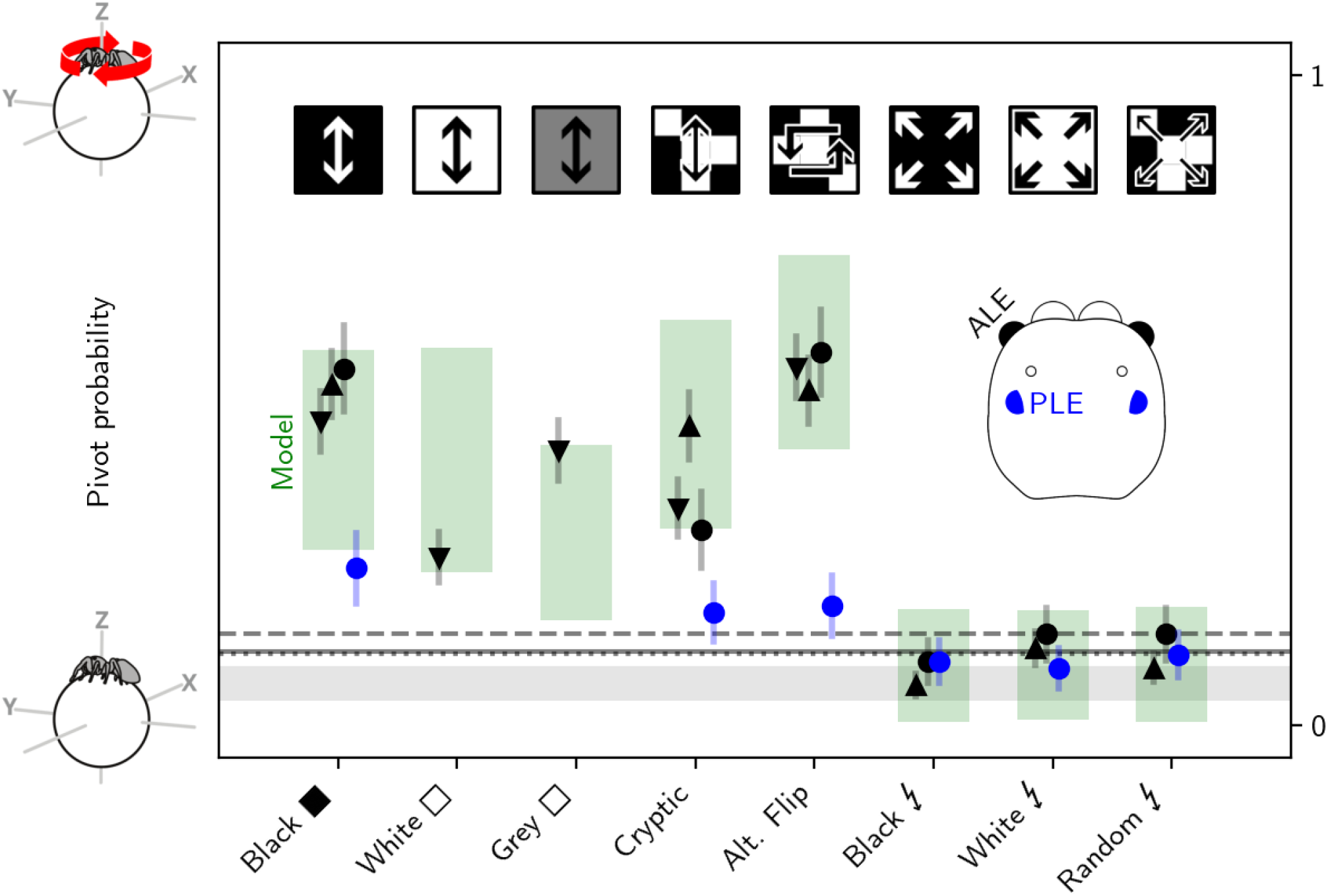
Spatiotemporal energy model predicts pivot behavior. Ordinate plots probability that the spider produces a pivot movement in response to different stimuli (abscissa; see also **Figure 1**) presented to different eye pairs (black for ALE, blue for PLE) during the course of three different experimental series (indicated by different symbols: down-triangle for series I, up-triangle for series II, and circle for series III). Horizontal lines indicate baseline response in the absence of any stimulus for different experimental series (solid for series I, dashed for series II, and dotted for series III). Shaded areas plot ranges (mean ± SD across repeated iterations) spanned by model simulations (green in response to stimulus, gray for baseline). Error bars show SEM.

**Figure 3:**
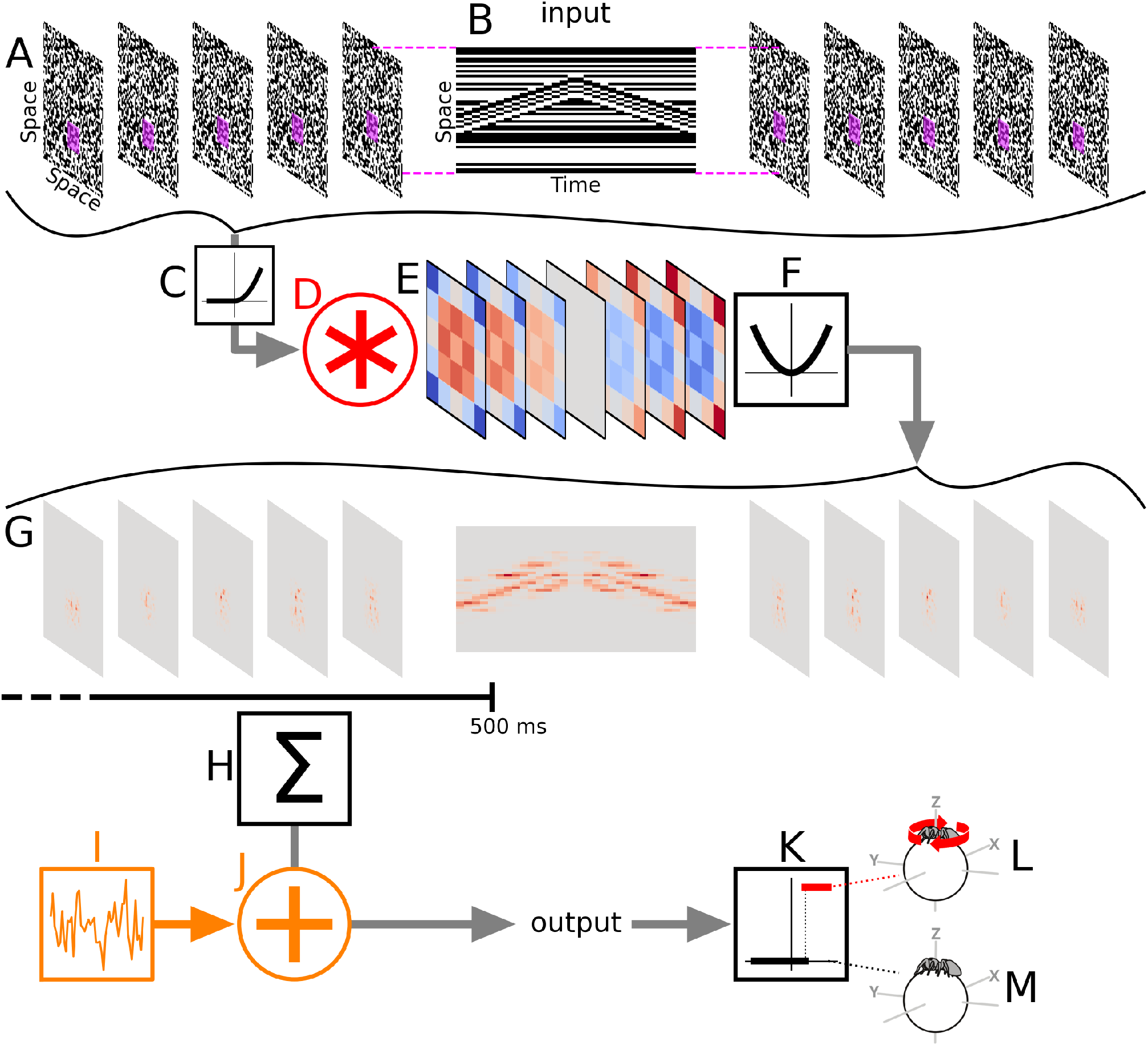
Model architecture. Input stimulus (**A**–**B**, plotted to the conventions of **Figure 1**) is subjected to a static expansive nonlinearity (**C**) before convolution (**D**) with a biphasic spatiotemporal filter (**E**, red/blue for positive/negative values) and subsequent squaring (**F**). The resulting output is plotted in G to align with input stimulus (red tint reflects response amplitude). This spatiotemporal output was converted to a scalar value by summing across space and over the first 500 ms of the stimulus (**H**). The resulting value was perturbed by an additive (**J**) Gaussian noise source (**I**), and thresholded (**K**) to generate a binary response corresponding to response (spider turns, **L**) or no response (spider remains still, **M**).

### 3.2 Experiment 2

In experiment 2, we removed the white and gray stimuli knowing that they can elicit a response. We instead added 3 new control stimuli. The “flash black” and “flash white” both present a segmentable square. This does not translate in space but rather appears and disappear. The “flash noise” instead both does not translate in space and does not present a segmentable square. Given the absence of white stimulus, in this experiment the subject selection was operated only on the black square. Seven out of the 36 subjects (19.4%) responded to less than 10% of the control stimuli, and as such were dropped from the subsequent analysis. We did observe a general effect of stimulus type (GLMM ANODA, *χ*^2^=393.53, DF=6, p<0.0001). Specifically, the spiders showed a response level significantly different from their baseline activity for the black, cryptic, and alternating flip stimuli (Figure 2. GLMM post-hoc, Bonferroni corrected. Black: odds ratio=8.991, SE=1.49, z=13.285, p<0.0001. Cryptic: odds ratio=0.144, SE=0.0236, z=11.809, p<0.0001. Alternating flip: odds ratio=0.116, SE=0.0191, z=13.066, p<0.0001.). The other three stimuli instead did not significantly differ from the no-stimulus control in term of response (Flash black: odds ratio=1.845, SE=0.484, z=-2.335, p=0.4108. Flash white: odds ratio=0.908, SE=0.19, z=0.459, p=1. Flash noise: odds ratio=1.267, SE=0.293, z=-1.024, p=1.). There was no difference between the first three stimuli nor between the other three.

### 3.3 Experiment 3

In experiment 3, the same stimuli of experiment 2 were presented. In this case however, the spider was placed between two separate computer monitors. The stimuli could appear either in an area visible to the ALEs or in an area visible to the PLEs, to check which type of stimulus elicits a reaction when perceived by the different eye-pairs. For this experiment, the subject selection was operated on the black stimuli presented in an ALEs-only position, as we have yet no expectation for a response rate to stimuli presented to PLEs. Eleven out of the 33 spiders (33.3%) were dropped from subsequent analysis. We did observe a significant effect of the stimulus type (GLMM ANODA, *χ*^2^=152.141, DF=6, p<0.0001), as well as an effect of the recruited eye-pair (*χ*^2^=35.239, DF=1, p<0.0001), and of the interaction between the two (*χ*^2^=16.518, DF=6, p=0.0112). The spider responded significantly different from the no-stimulus control for the black, cryptic and alternating flip stimuli, equally to experiment 2, and did so with both ALEs and PLEs (Figure 2. GLMM post-hoc, Bonferroni corrected. Black in ALE field: odds ratio=9.585, SE=2.53, z=8.564, p<0.0001. Black in PLE field: odds ratio=4.147, SE=1.24, z=4.756, p<0.0001. Cryptic in ALE field: odds ratio=3.383, SE=0.925, z=4.456, p=0.0002. Cryptic in PLE field: odds ratio=2.729, SE=0.877, z=3.124, p=0.0321. Alternating flip in ALE field: odds ratio=10.616, SE=2.82, z=8.907, p<0.0001. Alternating flip in PLE field: odds ratio=2.945, SE=0.932, z=3.411, p=0.0117). For the black and alternating flip stimuli, the response observed in the ALEs field was higher than the one observed in the PLEs field (Black ALE *vs*. PLE: odds ratio=3.825, SE=1.3, z=3.955, p=0.0014. Alternating flip ALE *vs*. PLE: odds ratio=5.966, SE=2.12, z=5.014, p<0.0001), while we observed no difference between the two eye-pairs for the cryptic stimulus (cryptic ALE *vs*. PLE: odds ratio=2.051, SE=0.753, z=1.958, p=0.905). No response different than the no-stimulus control was observed for the other three stimuli, equally to experiment 2, with neither ALEs or PLEs (Flash black in ALE field: odds ratio=0.855, SE=0.314, z=-0.426, p=1. Flash black in PLE field: odds ratio=1.415, SE=0.535, z=0.918, p=1. Flash white in ALE field: odds ratio=1.285, SE=0.42, z=0.767, p=1. Flash white in PLE field: odds ratio=1.252, SE=0.491, z=0.572, p=1. Alternating flip in ALE field: odds ratio=1.285, SE=0.42, z=0.767, p=1. Alternating flip in PLE field: odds ratio=1.584, SE=0.58, z=1.256, p=1).

### 3.4 A simple computational model captures all results

Because our stimuli span a wide range of specifications, it is difficult to interpret the associated behavioral responses on the basis of intuitive considerations about stimulus content (e.g. whether the presented pattern contained movement or not, or whether it contained a homogeneous versus heterogeneous region of interest). To aid our interpretation, we implemented an explicit computational model of spider behavior that borrows concepts from existing architectures used to capture visual processes in insect vision [41, 42].

The structure of our model is simple (see **Figure 3**): after front-end nonlinear transduction, each input stimulus is convolved with a spatiotemporal energy filter (**Figure 3E–F**). The resulting output (an example is plotted in **Figure 3G**) is briefly integrated over an initial temporal window of half a second to generate the decision variable, which is then thresholded to produce a binary decision (pivot versus stay, **L** versus **M**). The decision variable is perturbed by an internal noise source (**I**) to capture behavioral variability. As shown in **Figure 2** (shaded regions), this model captures all our empirical observations. It is notable in this respect that the model does not incorporate any directional element: it does not support motion direction discrimination.

We regard our model as minimal in the following sense. In developing the model, we proceeded by adding elements only when dictated by data. For example, we started with a monophasic spatiotemporal filter (in place of the biphasic filter shown in **Figure 3E**), but we found that this filter specification failed to reproduce spider responses to certain stimuli, prompting us to modify the filter temporal profile. The same process applied to the addition of pre-/post-filter nonlinearities (**C**/**F**). For example, in the absence of the front-end nonlinearity (**C**), the model would capture most of the behavior in **Figure 2** except for the amplitude of the response to the grey-square stimulus (third stimulus on the x axis in **Figure 2**), motivating the addition of this element to the model. Although we cannot exclude that simpler architectures may produce comparable results, we think it unlikely that our model can be substantially simplified from its current implementation.

## 4 Discussion

In this study, we tested the process of detection of moving target by jumping spiders using their secondary eyes, and which characteristics govern their pivot behavior. We observed that the spiders produced pivot turning towards any spatio-temporal changes in luminance rather than to the presence of segmentable targets. On the other hand, they did not respond to flashes—luminance changes that, while some even presented clear objects, did not move spatially. We also observed that the detection probability is spread across all secondary eyes, with no specific pair being specialized for any type of motion. Finally, we developed a spatiotemporal filter describing the spiders’ behavior.

### 4.1 The spiders’ behavior can be explained without a motion detector

The results of this experiment were surprising. Jumping spiders are quite effective visual predators, and they can also be quite selective about the types of visual targets they turn towards or focus on [22, 17, 18, 16]. Accordingly, we expected that the secondary eyes would induce the production of pivot upon the detection of real objects characterized by definable contours and a coherent motion trajectory. Crucially, instead, the spiders produced pivots towards the Alternating Flip, which does not constitute a segmentable object nor is technically in motion. In psychophysical terms in fact, motion detection refers to the decoding of a translation direction of the objects in the visual field, rather than the detection of any change in luminance [48]. In this sense, the Alternating Flip stimulus does not constitute a moving stimulus, but only an example of spatio-temporal luminance change. These results would suggest that motion perception is not useful in the process of target detection for jumping spiders, that instead solves the task with filters only.

The idea that jumping spiders may not possess any circuit for motion detection is provocative. The presence of these structures is well established across the evolutionary tree, formally modeled in its most simple form as the Reichard-detector [48] and even fully identified at the neuronal level in *Drosophila* [49]. At least in their behavior, jumping spiders seemed to perfectly match the same computational pattern. We know from previous literature, in fact, that these animals are particularly attracted to specific object sizes and motion speeds [12, 14], are extremely sensitive to specific motion directions, such as looming objects [20], and can even discriminate between complex motion patterns even when their low-level characteristics are matched (as is the case for biological motion point-light displays, [22, 13]). Yet, for none of these tasks do either the target position or its translation direction strictly need to be encoded. Simple filters, like the one developed for this paper, are sufficient to govern the jumping spider pivot response in all of the aforementioned cases. Even the complex biological motion patterns may be decoded, in theory, by simply computing local coherency [13]. Jumping spiders even seem to be immune to at least some motion illusion [50], the perception of which actually rests on the functioning of the motion detection circuit [51].

In a previous study, we tested *Menemerus semilimbatus* spiders in an Habituation/Dishabituation paradigm, using motion direction as the varying stimulus characteristic in the dishabituation presentation [52]. There, the animal successfully increased the response rate upon the presentation of the dis-habituation stimulus, demonstrating they could decode the difference between targets. Yet, the stimuli were not controlled for the area of the visual field occupied, making the task solvable even without directly encoding for the motion direction.

To date, no direct evidence is available demonstrating that jumping spiders are actually capable of motion detection. If this is the case, these animals would represent a fascinatingly unique example across developed visual systems. The visual behavior of jumping spiders remains in fact deeply complex even if deployed in a nervous system the size of a poppy seed. Doing away without motion perception circuit would represent a bold example of neural simplification, reducing cognitive load while apparently maintaining function. This hypothesis should, however, be thoroughly tested, especially given its uniqueness.

### 4.2 Lack of response to the flashing stimuli

In none of the experiments, the animals reacted to any of the flashing stimuli, being them a full object (like in the black and white flashes) or scattered dots (like in the noise flash stimulus). It is crucial to point out however that a lack of response in our experimental paradigm does not necessary suggests that the spiders failed in the detection of the stimuli. As stated above, in a previous experiment we already used stimuli similar to the flashing objects here presented [19], and already observed a lack of response from the spiders. Yet, the presentation of such stimuli caused the spiders to change both reaction time and probability for the presentation of a successive translating dot. This demonstrate that the spiders did detect the flashes, even if these did not elicit a pivot response.

It is in our opinion most likely that the spatio-temporal filter presented in this paper is implemented relatively late in the visual hierarchy of the jumping spiders nervous system, the results of which is then threshold to determine the production of a pivot response (see Figure 3). Other pathways may use the detected luminance changes in the visual scene in a variety of different tasks, like the allocation of attention [19] without eyes of body shifting. Given the setup of the presented experiments we cannot determine if this is the case, as the pivot response is the only behavioral measure available. The detection of early neural responses should probably be inquired upon using electrophysiological techniques.

### 4.3 Difference between eyes

In experiment 3, we tested the differential ability of ALEs and PLEs to detect and react to the various type of moving stimuli. We did so by presenting the stimuli in areas of the visual field only visible to ALEs and PLEs respectively, identified accordingly to the visual spans presented in [13]. The reaction probability observed for ALEs are consistent with the results of experiment 1 and 2, congruently with the fact that in these two cases the stimuli were presented frontally to the spiders, and as such were already implicitly visible to the ALEs only. The spiders’ response pattern to the PLEs’ presented stimuli was roughly the same as the ALEs’ one, exception made for a lower overall response probability. The latter is probably to be linked to the general lower visual acuity of the PLEs in respect to ALEs, which in turn would make the stimuli presented in these sections of the visual field harder to detect.

The existing literature is generally concordant in stating that both ALEs and PLEs are responsible for motion detection [8]. Yet, there is an increasing amount of evidence that suggests that ALEs only are specialized in the finer motion discrimination tasks [22, 13, 20, 14], making them somewhat unique among all secondary eyes. Accordingly, we would have expected the spiders to respond to the first-order stimuli with both ALEs and PLEs, and instead to the second-order stimuli only when presented with the ALEs, most likely the only pair equipped with the specific motion-detection circuit required for these type of stimuli. Yet as already stated, the spiders’ pivots are triggered without any specialized motion-detection circuit, making the differences between ALEs and PLEs irrelevant for the task: in the presence of a single stimulus, the finer ability of ALEs to discriminate types of motion is irrelevant, as the spiders will nonetheless be pushed to perform a pivotal rotation to fixate the stimulus with AMEs.

Regardless, observing these responses to be spread across secondary eyes, together with the segregated structure of the jumping spiders’ visual field [53], allow us to make inferences on the most likely candidate brain region to implement the spatio-temporal filter, as it necessarily need to be shared by ALEs and PLEs. We suggest this to be the arcuate body (AB). This area has previously been observed to have a retinotopical structure [54, 55], likely spread across the spiders’ combined visual field given it receives connections from all eyes [53]. Moreover, the AB presents a layered, columnar structure [56, 53], which would be more than capable of supporting a convolution of the visual field across space and time.

### 4.4 Conclusions

In this paper, we tested the ability of jumping spiders to detect moving objects with increasingly more complex luminance patterns. Using a psychophysical model, we demonstrated that the spiders can solve the task using a single spatio-temporal filtering process, rather than relying on a different computational process for each stimulus type as we previously believed. What is more, by behaviorally identifying the spread of the detection ability across eyes, we infer the most likely brain region where the modelled filter is deployed. Our study opens up on the possibility that the visual computational structure of jumping spiders follows a different logic from all other animals, from vertebrates and invertebrates alike, doing away with widespread circuitry in favor of simpler solution, an example of efficiency in the process of brain miniaturization.

